# Evaluating Invasive EEG Implantations in Medically Refractory Epilepsy with Functional Scalp EEG Recordings and Structural Imaging Data

**DOI:** 10.1101/554667

**Authors:** Anil Palepu, Adam Li, Zachary Fitzgerald, Katherine Hu, Julia Costacurta, Juan Bulacio, Jorge Martinez-Gonzalez, Sridevi V. Sarma

## Abstract

Seizures in patients with medically refractory epilepsy (MRE) epilepsy cannot be controlled with drugs. For focal MRE, seizures originate in the epileptogenic zone (EZ), which is the minimum amount of cortex that must be treated to be seizure free. Localizing the EZ is often a laborious process wherein clinicians first inspect scalp EEG recordings during several seizure events, and then formulate an implantation plan for subsequent invasive monitoring. The goal of implantation is to place electrodes into the brain region covering the EZ. Then, during invasive monitoring, clinicians visually inspect intracranial EEG recordings to more precisely localize the EZ. The EZ is then surgically removed. Unfortunately surgical success rates average at 50%. Such grim outcomes call for analytical assistance in creating more accurate implantation plans from scalp EEG. In this paper, we introduce a method that combines imaging data (CT and MRI scans) with scalp EEG to derive an implantation distribution. Specifically, scalp EEG data recorded over a seizure event is converted into a time-gamma frequency map, which is then processed to derive a spectrally annotated implantation distribution (SAID). The SAID represents a distribution of gamma power in each of the eight cortical lobe/hemisphere partitions. We applied this method to 4 MRE patients who underwent treatment, and found that the SAID distribution overlapped more with clinical implantations in success cases than in failed cases. These preliminary findings suggest that the SAID may help in improving EZ localization accuracy and surgical outcomes.

## I. Introduction

Over 60 million people worldwide have epilepsy, and approximately 30% have medically refractory epilepsy (MRE) in that their seizures cannot be controlled by drugs [4], [5], [8], [24]. For focal MRE patients, seizures originate in the epileptogenic zone (EZ), which is the minimum amount of cortex that needs to be treated in order to eliminate seizures [27], [33]. There are currently three treatments for focal MRE: i) surgical resection of the EZ, ii) laser ablation of the EZ, or iii) stimulation of the EZ. In order for any of these treatments to work, clinicians must successfully localize the EZ.

The localization process entails (i) obtaining scalp EEG recordings from a patient over several seizure events, (ii) formulating a hypothesis of where the EZ is (both in hemisphere, lobe and/or structures), and then if required, (iii) implanting electrodes either at the surface of the cortex or deep in the brain to cover the hypothetical EZ region. Clinicians then visually inspect the intracranial EEG (iEEG) recordings over several seizure events to precisely localize the EZ. Despite this lengthy monitoring process and despite the fact that large brain regions may be removed, surgical success rates vary between 30%-70% [9], [20], [29], [41], [44].

Such variable outcomes are often due to challenging cases where no lesions appear on MRI scans. In these cases especially, failed treatment may be partially attributed to incorrect or insufficient invasive electrode coverage. With the boom of big data and increased computational power, computational models should be used to assist clinicians in forming an implantation plan from scalp EEG recordings. There are many algorithms that operate on scalp EEG to help localize the EZ [1], [2], [35], [38]. They directly try to predict the region of the EZ. However, scalp EEG is insufficient to localize deeper structures that may be involved in seizure events [32]. These approaches do not directly align with how clinicians currently use scalp EEG, which is to form an implantation hypothesis (i.e. which lobes to insert iEEG electrodes) for subsequent invasive monitoring.

In this study, we develop an algorithmic approach to assist clinicians in forming an implantation hypothesis from scalp EEG. Our method processes a patient’s scalp EEG recordings over seizure events in addition to structural data from the patient’s T1 MRI and CT scans to generate an implantation distribution. More specifically, scalp EEG recordings are converted into a temporal-gamma band map, which is then converted into a spectrally annotated implantation distribution (SAID). The SAID quantifies the power in the gamma band over a seizure event in each of eight hempishere-lobe partitions (HLPs) of the brain: Left/Right Frontal (LF, RF), Left/Right Parietal (LP, RP), Left/Right Occipital (LO, RO), Left/Right Temporal (LT, RT).

We applied our method to data captured from 4 MRE patients treated at the Cleveland Clinic, and each SAID was compared to the CAID by computing a distance between the two distributions (see fig. 1). We found that the SAID and CAID are more similar in patients who had successful surgical outcomes and less similar in patients who had failed surgical outcomes. These preliminary findings suggest that gamma rhythms in EEG may be correlated to seizure onset activity, and thus helpful in localization of the EZ in focal MRE.

**Fig. 1:**
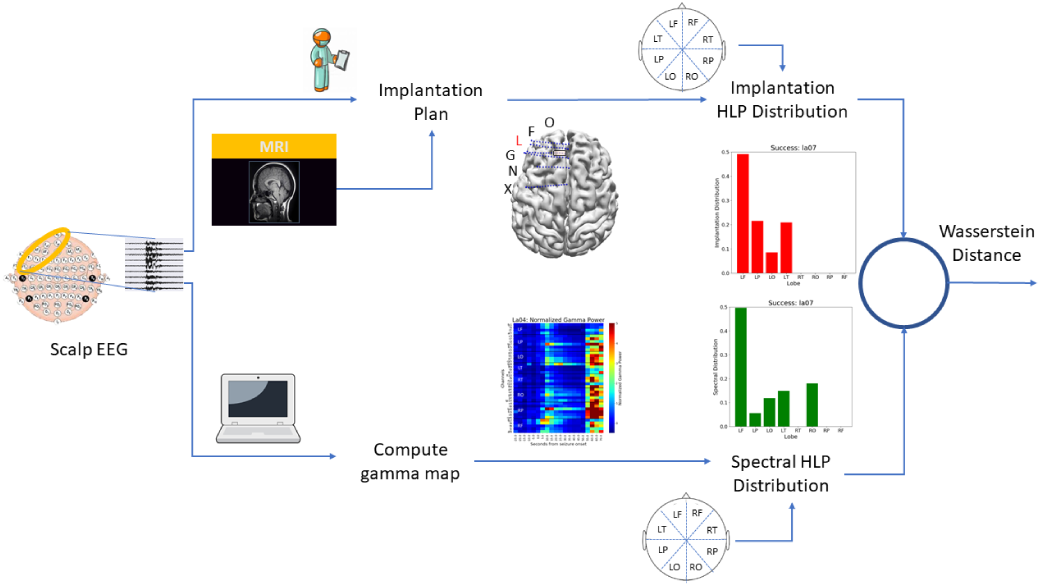
Analysis pipeline for generating SAID and CAID.

## II. METHODS

### A. Epilepsy Patient Data - Scalp EEG, T1 MRI, CT

The patients included in this study were surgically treated for drug-resistant seizures at the Cleveland Clinic (CC). All underwent invasive presurgical monitoring with depth electrodes for seizure localization or mapping of eloquent areas. Scalp EEG data was collected in conjunction with video EEG.

The SEEG technique uses several small drill holes (1.8 mm in diameter), allowing many electrodes to be inserted (up to 20). SEEG may provide a more complete coverage of the brain, from lateral, intermediate and/or deep structures in a three-dimensional arrangement recorded over hundreds of channels. The number and location of implanted electrodes are pre-operatively planned based on a pre-implantation hypothesis, which is formulated in accordance with non-invasive pre-implantation data such as seizure semiology, ictal and interictal scalp EEG, MRI images, PET and ictal SPECT scans. Intracranial contact locations were documented by postoperative CT coregistration with preoperative MRI. Decisions regarding the need for invasive monitoring and the placement of electrode arrays were made independently of this work and solely based on clinical necessity. The research protocol was reviewed by the Institutional Review Board (IRB) at the CC.

Pre-operative MRI scans (T1, contrasted with Multihance, 0.1mmol *kg*^-1^) were obtained prior to electrode implantation for use in the electrode placement procedure. CT scans were performed after electrode implantation in order to ensure accurate placement and labeling. Following implantation surgery, a three-dimension reconstruction was produced and visualized from coregistered MRI and CT scans in Curry NuroImaging Suite (NeuroScan, El Paso, TX) and the location of each electrode determined with visual inspection and agreement by at least two clinical experts. The data for this study was obtained through retrospective review and stored on an IRB-approved encrypted hard drive. T1 and CT data was processed using FreeSurfer, FSL and Fieldtrip Toolbox [16], [21], [31].

Scalp EEG recordings were acquired for clinical purposes independent of this study. The recordings were initially acquired with Nihon Kohden system (Nihon Kohden Corp.) with 200Hz sampling rate, 300Hz antialiasing filter, and archived on the IRB-approved Epilepsy Monitoring server. Data were then retrospectively obtained for review and converted to EDF format for deidentification and assigned study numbers (e.g. la05), which were then given to us for processing in no specific order. Digitized data were stored on an encrypted IRB-approved hard drive compliant with Health Insurance Portability and Accountability (HIPAA) regulations. Board-certified electroencephalographers marked, by consensus, the unequivocal electrographic onset of each seizure and the period between seizure onset and termination. Data was then preprocessed as .fif and .json files for analysis in Python with data I/O facilitated by MNE-Python [18], [19].

### B. Computing the Spectral Annotated Implantation Distribution (SAID) from Scalp EEG

All data underwent digital filtering with a butterworth notch filter of order 4, implemented in SciPy with frequency ranges of 59.5 to 60.5. We applied a common average referencing scheme to the data before analysis [28]. This has been shown to produce more stable results and rejects correlated noise across many electrodes [17]. We made sure to exclude any electrodes from subsequent analysis if they were informed to have artifacts in their recording by clinicians. We then considered the discrete fourier transform (DFT) (eq 1), which breaks down a electrode signal, *x*_*n*_ with n as the index through samples of the signal, into its sinusoidal components with certain magnitude and phase [7].

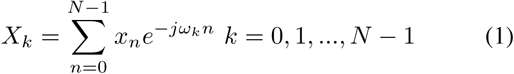

This generates the relative contribution of each frequency component at each point in time for every electrode. We then represent each electrode’s DFT in terms of its frequency bands, where we defined the gamma band as 30-100 Hz. We average the power for all frequencies in the gamma band for each electrode into a gamma map, *G*. Then for each electrode and its gamma band, we select the baseline data as the preictal data for that patient. We take a selected window of 10-25 seconds after clinically annotated seizure onset ([*W*_1_, *W*_2_]) and compute the shift versus the preictal baseline. Finally, we normalize the computed gamma power in this time window by subtracting the mean and dividing by the standard deviation of the gamma power in each corresponding electrode of the baseline data to compute 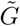 (see eq 2).

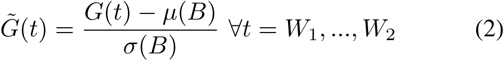

In the scalp EEG, we can discretize the contacts into eight regions, or eight HLP lobes of the brain for left and right hemispheres: Frontal (LF, RF), Temporal (LT, RT), Parietal (LP, RP) and Occipital (LO, RO) lobes. It is worth noting again that scalp EEG is insufficient to record signals from deeper structures, such as the limbic, or insular regions [32]. So we mainly focus on the cortical surface lobe partitions of the brain. From this data, we can compute an HLP distribution by taking the mean normalized gamma power across all electrodes and selected times in each lobe to calculate the mean power of each lobe in the selected window. These 8 mean power values together will form the SAID for a specific patient.

### C. Computing the Clinical Annotated Implantation Distribution (CAID) of SEEG Contacts

Given the SAID, we aim to compare it with the ground truth of clinical implantations. In order to localize the iEEG contacts in T1 MRI brain space, we first localize the contacts in their CT image, apply an affine transformation to map these coordinates into T1 space, then apply a segmentation algorithm to map each voxel in the T1 image to a brain region, and finally apply a dictionary mapping of each brain region to a lobe (see fig. 2). We localize the contacts in the CT image using open-source software from the Fieldtrip Toolbox [31]. Contacts with their corresponding xyz coordinates in the CT image volume are obtained and then mapped to the T1 MRI image volume space via an affine transformation computed by the open-source software FSL [21], [22].

**Fig. 2:**
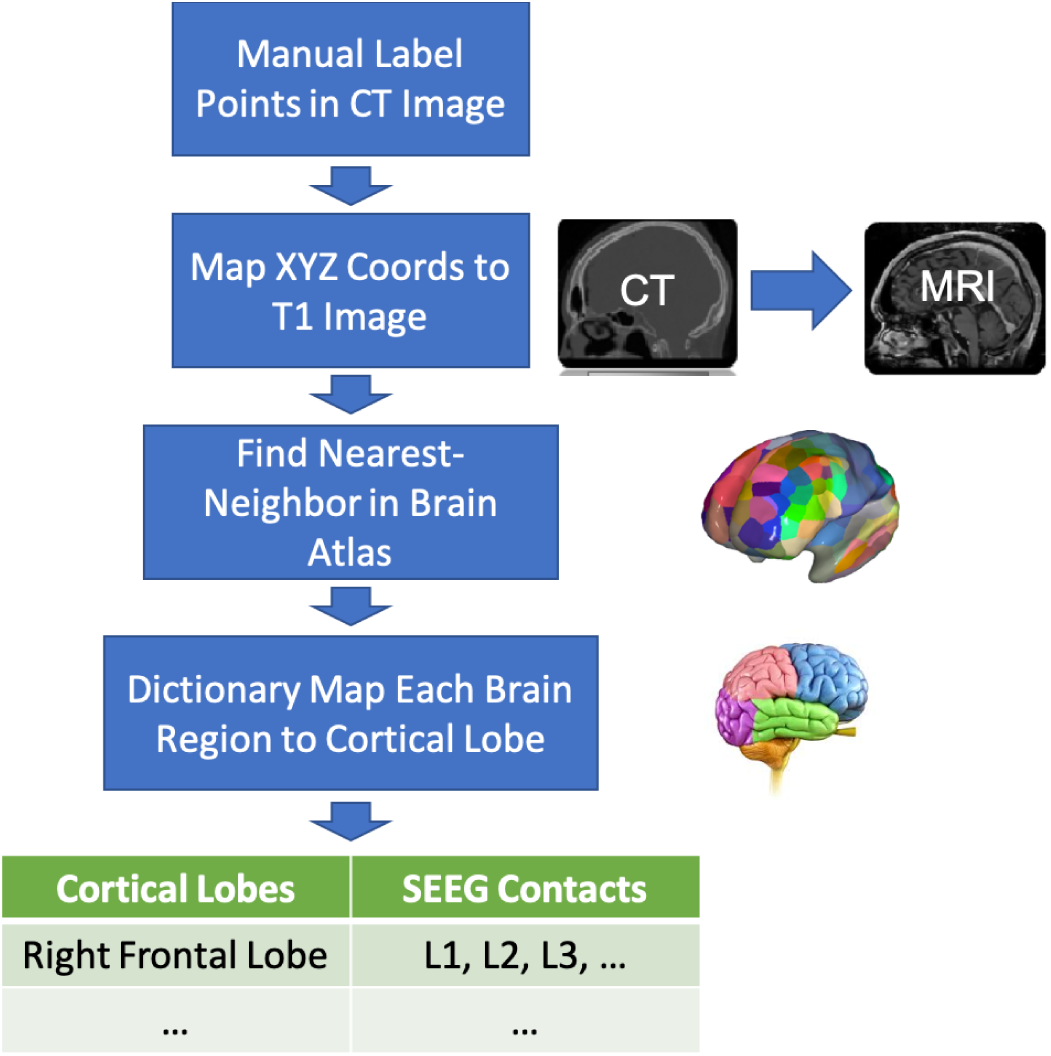
An outline of how the clinical HLP distribution is computed in a quantitative process.

It is worth noting that there are other available methods of neuroimage coregistration, such as robust registration and image large deformation diffeomorphic metric mapping [3], [37]. These methods are all able to be used during the coregistration portion of our described procedure. In order to make sense of the SEEG coordinates in the T1 MRI brain space, we perform automated segmentation and parcellation of the T1 MRI image volume using FreeSurfer [11], [15], [16]. This segments the brain into either the Desikan-Killany (34 cortical regions per hemisphere), or the Destrieux atlas (74 cortical regions per hemisphere) [12], [13], which assigns each voxel to an annotated brain region. We then apply a dictionary mapping that assigns each of these unique brain regions to a specific lobe: Frontal (F), Temporal (T), Parietal (P), Occipital (O), Limbic (L), and Insular (I). Given the centroid coordinates of each atlas brain region, we apply a nearest-neighbor algorithm to assign each SEEG contact to its nearest brain region. Then, we use the dictionary mapping of these regions to assign the contact to a specific brain lobe. Depending on the contact’s hemisphere, if a result is in the Frontal, Temporal, Parietal, or Occipital lobes, we assign the contact to its corresponding HLP lobe. Results in the Limbic or Insular lobes are omitted, as they are not easily assigned to 1 of the 8 HLP lobes. These 8 hemisphere/lobe SEEG contact assignments form the CAID for the patient.

### D. Comparing SAID to CAID

In order to quantitatively compare the SAID to the CAID, we make use of the Wasserstein metric in eq 3, which computes a distance metric between the distributions *u* and *v*, where Γ(*u, v*) is the set of distributions on ℝ *×* ℝ. We compute *W* for each patient’s CAID vs SAID and summarize the distances for successful and failed surgical patients. An example of CAID and SAID for two patients is shown in fig. 4a and 4b with the resulting Wasserstein distances shown in fig. 4c.

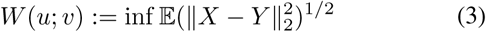

Our code that describes this entire pipeline is available online at: https://github.com/adam2392/neuroimg_pipeline. In addition to code made open-source, we contribute a Docker and Singularity container to allow easy usage of the software to compute CAID and SAID.

## III. Results

In this section, we show results of our proposed methodology applied to four patients with invasive SEEG implanted. We apply the DFT algorithm to sliding windows of data around the seizure onset, and compute the power within the gamma band. In fig. 3, we show an example of the gamma power around seizure onset that is normalized against a preictal baseline computed from each patient’s scalp EEG data. As seen in other studies, we expect a baseline shift in gamma power that is related to a seizure starting [36], [43].

**Fig. 3:**
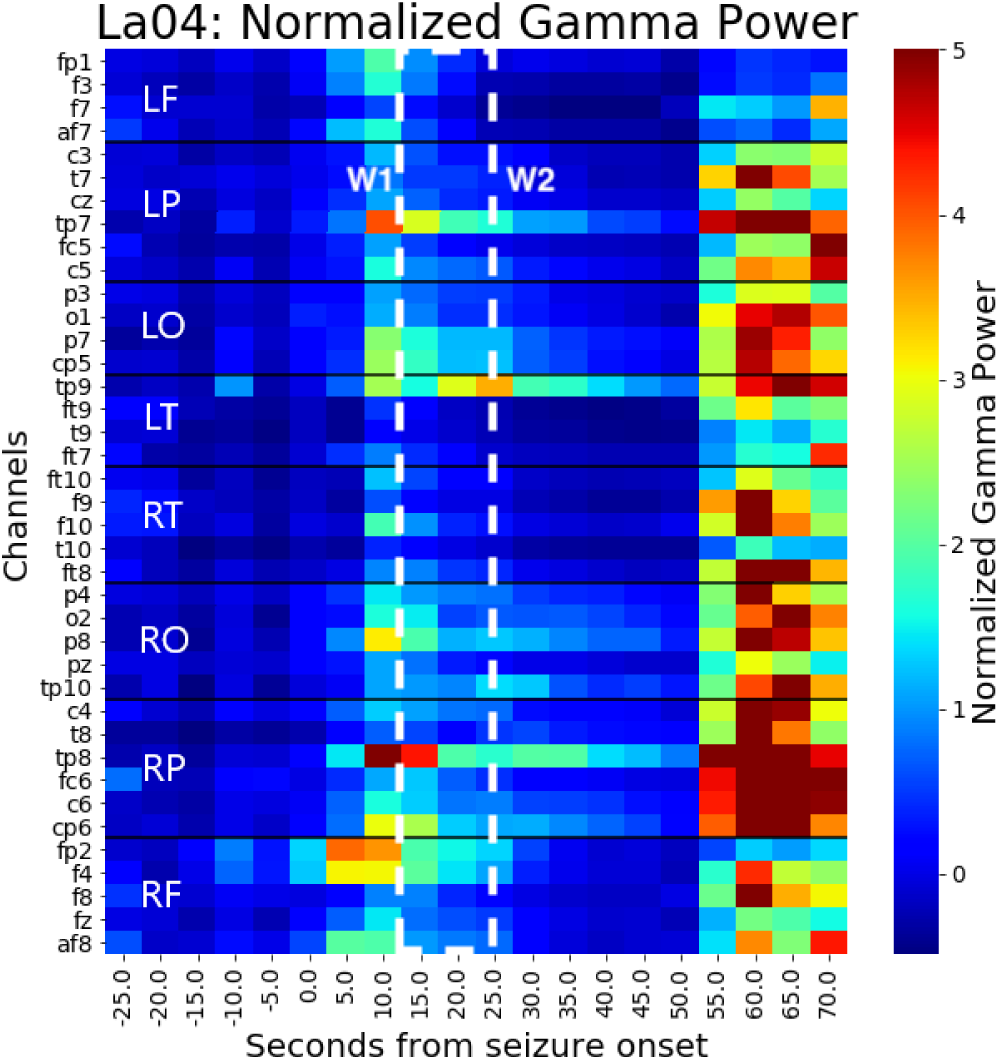
An example of the normalized gamma power for patient identifier LA04, with left-frontal lobe epilepsy. The red indicates a relative increase from baseline gamma power, while blue indicates power more similar to the baseline. The channels are organize into their respective right and left lobes. The dashed lines inside this figure show the selected windows of time used to compute 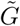 in eq. 2.

We then compute the corresponding HLP distributions for each patient and compare them using the Wasserstein distance. In fig. 4, the CAID is more similar to the SAID for successful outcomes. In two of the successful patients, we see a lower Wasserstein distance between the SAID and CAID, while we see a higher distance in failure patients.

**Fig. 4:**
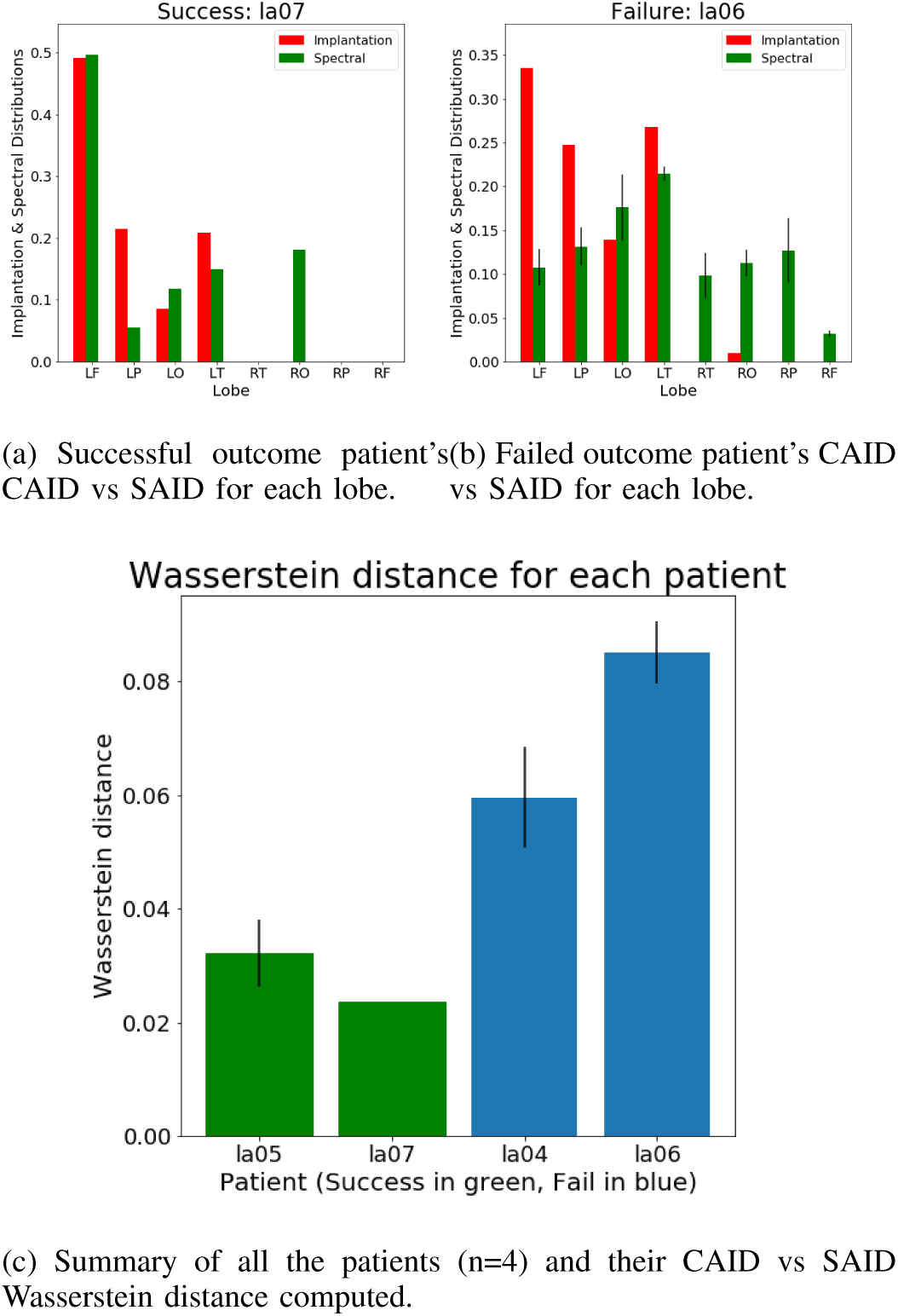
This figure shows an example of a successful (a) and failed (b) surgical outcome patient’s CAID vs SAID distributions. Then we show the computed Wasserstein distance metric between CAID and SAID for each of the four patients analyzed. Note that error bars are standard error of the mean plotted for patients with multiple data snapshots of seizure events. For patient LA07, only one seizure event was provided, so there are no error bars for this patient.

## IV. Discussion

In this study, we examine a quantitative method for comparing spectral power in scalp EEG to clinical iEEG implantations. We do this by defining the contact locations in brain space for a patient through their T1 MRI and CT data. Applying a spectral model to the data, we can then compute the spatiotemporal dynamics of the gamma power band. Both the iEEG implantations and spectral power are able to be partitioned into eight cortical lobes, which form the CAID and SAID. We demonstrate that the SAID could potentially be used as a metric for planning implantation distributions. In our preliminary study on four patients, we show that the CAID is more similar to the SAID in successful surgical outcomes, while being more dissimilar in failed surgical outcomes.

Although the method we propose is quantitative, there are errors introduced along each step of the process that could affect results. Firstly, scalp EEG is insufficient to localize insular, or limbic regions as these are deeper structures, which can not be directly recorded from. A potential way to augment this approach is with diffusion-weighted MRI, which can give insight into the structural connections of the brain. In future work, we are interested in using this data to infer possible necessary implantations into the insular and limbic regions of the brain. In the T1 segmetation process, the process maps brains onto a set of presegmented and annotated atlases. This introduces errors into the process, as every brain is not the same. In addition, when the contact coordinates are localized, the accuracy depends on the resolution of the CT scans, which differ per patient. In addition, a coregistration algorithm is never perfect and will introduce errors in exactly where the coordinates are in brain space (i.e. T1 image). Furthermore, the seizure onset time marked by clinicians is done visually, meaning it can occur well before ictal activity has propogated to the scalp level and can be measured by the scalp EEG, especially in cases where the seizure onset is deep below the cortical surface. Because a fixed window around seizure onset is always used, the characteristics of the data used to compute power in the gamma band could vary between different patients and seizure events depending on this time delay.

Our framework can be extended further by allowing flexibility in the choice of brain atlas [6], [13], [14], [39], segmentation algorithm [30], [40], [42], and coregistration algorithm [3], [21], [22]. In addition, this framework is not limited to a spectral model of the data. In theory, any model that is applied on the scalp EEG contacts can be used, such as models that include other frequency bands [1], [2], [10], [45], measures of stability [25], personalized whole-brain models [23], [26], [34], or measures of graph statistics [10]. Future work will include analyzing a larger cohort of patients and a suite of different models to rigorously evaluate invasive EEG implantations in MRE epilepsy patients.

## Acknowledgment

AL is supported by NIH T32 EB003383, NSF GRFP, Whitaker Fellowship and the Chateaubriand Fellowship, and SVS is supported by NIH R21,NS103113.

